## Supplementary Material for "Metabolic landscape of the tumor microenvironment"

### Supplementary Figure 1

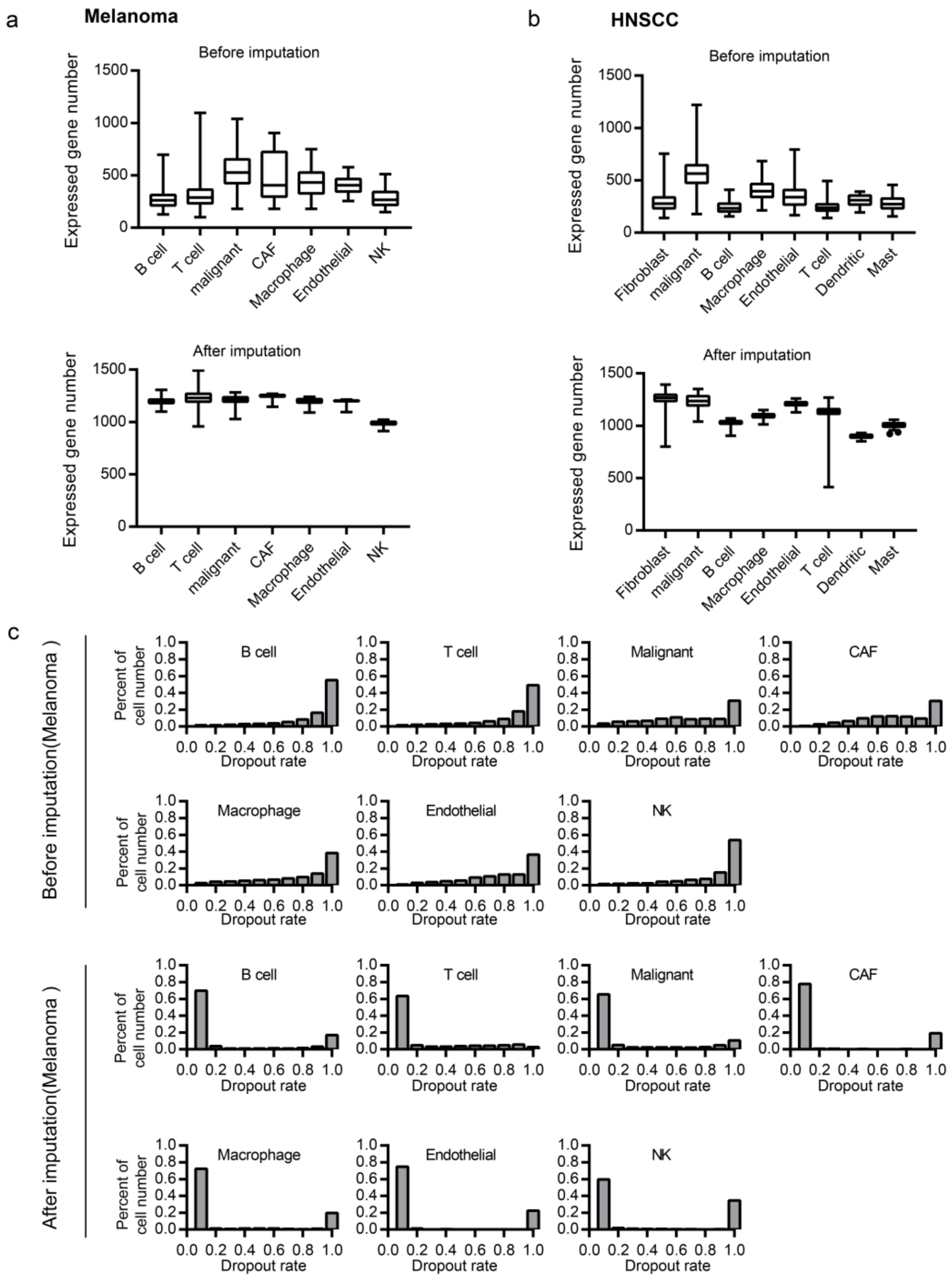

### Supplementary Figure 1 (continued)

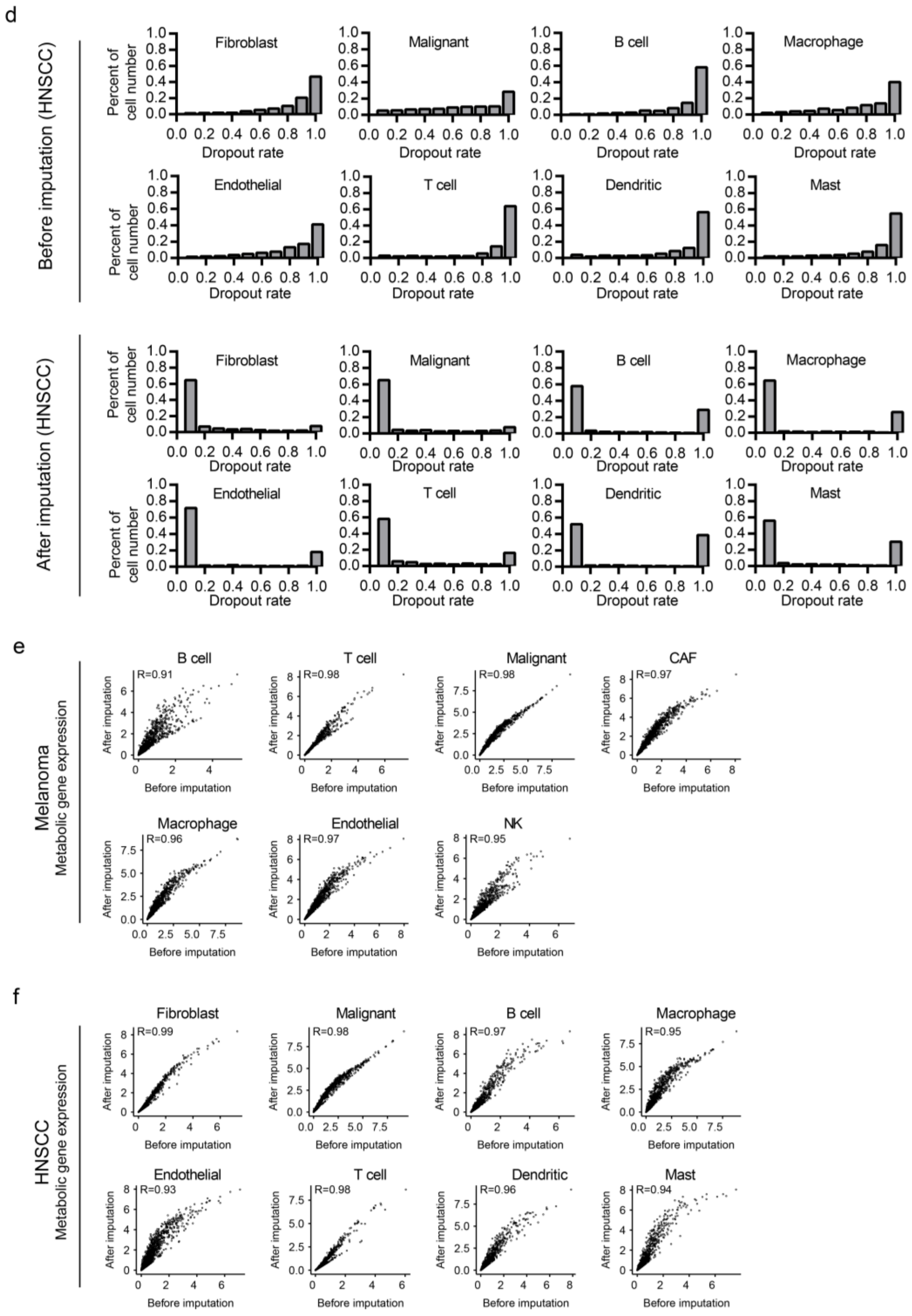

**Supplementary Figure 1. (Related to Figure 1)**

(a) Distributions of numbers of expressed genes (i.e. genes with non-zero expression value) before and after imputation in different cell types from the melanoma dataset.

(b) Same as in (a) but for the squama cell carcinoma of head and neck (HNSCC) dataset.

(c) Distributions of drop-out rates (i.e. fraction of zero gene expression values) in different cell types before and after imputation in the melanoma dataset.

(d) Same as in (c) but for the HNSCC dataset.

(e) Scatter plots comparing average metabolic gene expression levels before and after imputation in different cell types from the melanoma dataset.

(f) Same as in (e) but for the HNSCC dataset.

### Supplementary Figure 2

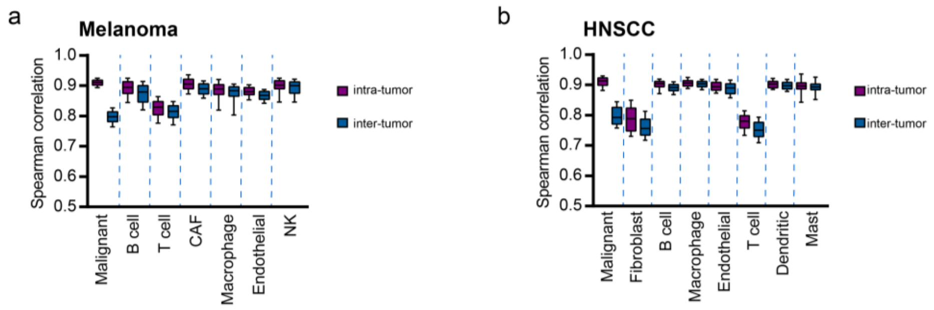


**Supplementary Figure 2. (Related to Figure 1)**

(a) Distributions of Spearman's rank correlation coefficients of metabolic gene expression between 500 randomly selected pairs of cells from the same tumor (i.e. intra-tumor) or from different tumors (i.e. inter-tumor) in the melanoma dataset.

(b) Same as in (a) but for the HNSCC dataset.

**Supplementary Figure 3**

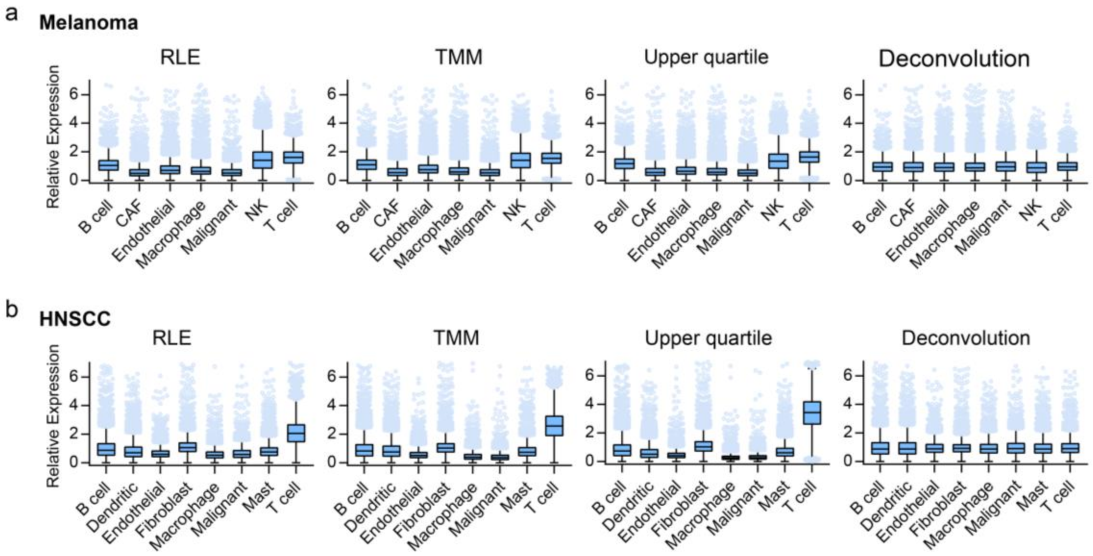


**Supplementary Figure 3. (Related to Figure 2)**

(a) Distributions of relative gene expression levels in cell types from the melanoma dataset after data normalization using relative log expression (RLE), trimmed mean of M-values (TMM), upper quartile or deconvolution method.

(b) Same as in (a) but for the HNSCC dataset.

### Supplementary Figure 4

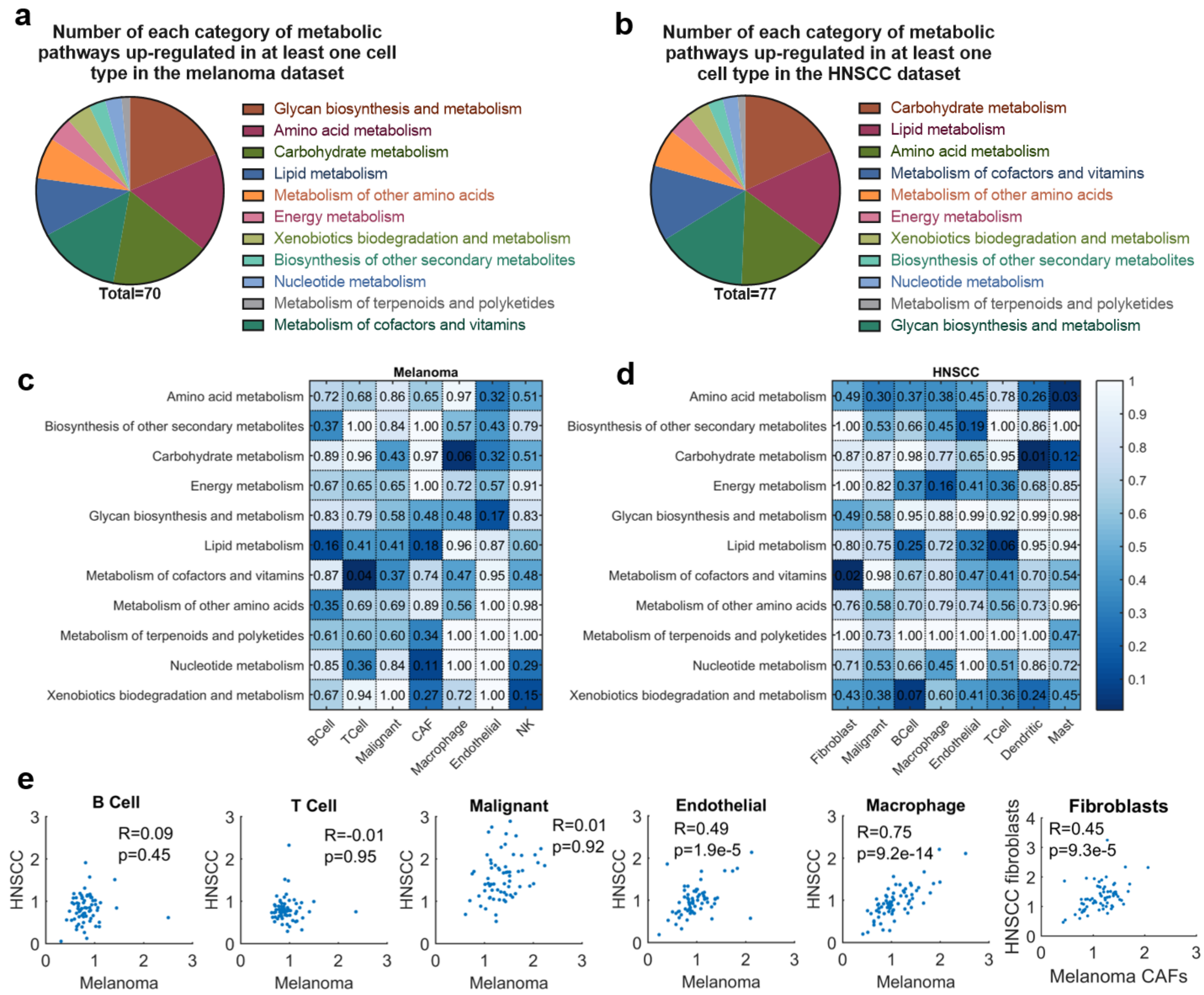


**Supplementary Figure 4. (Related to Figure 2)**

(a) Numbers of different categories of metabolic pathways up-regulated in at least one cell type in the melanoma dataset.

(b) Same as in (a) but for the HNSCC dataset.

(c) One-sided Fisher's exact test p-values for enrichment of categories of pathways in pathways up-regulated in different cell types in the melanoma dataset.

(d) Same as in (c) but for the HNSCC dataset.

(e) Scatter plots comparing metabolic pathway activities between the melanoma and HNSCC datasets for cell types shared by the two datasets.

### Supplementary Figure 5

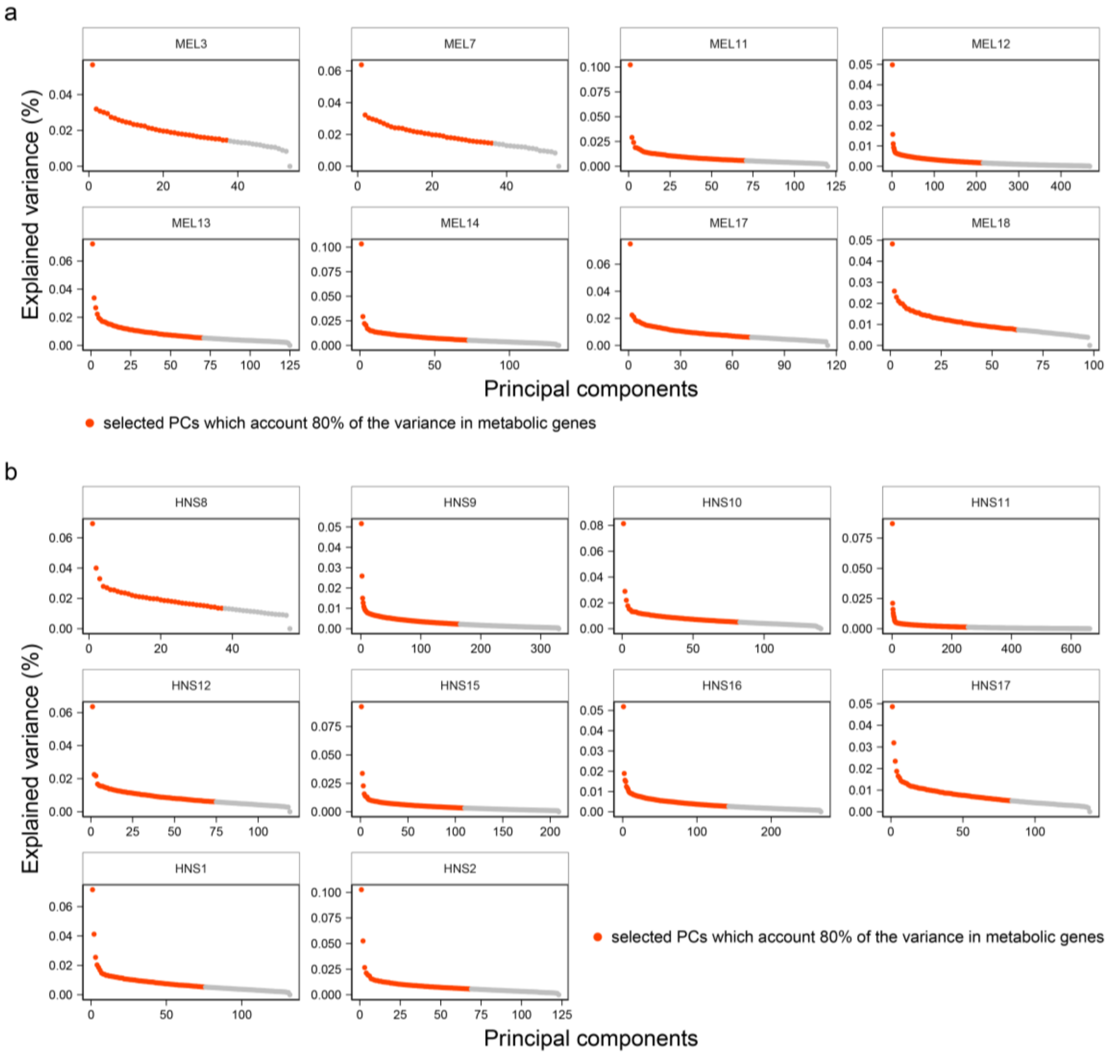


### Supplementary Figure 5. (Related to Figure 3)

(a) Explained variance of principal components (PCs) from principal component analysis (PCA) of metabolic gene expression levels in malignant cells from different tumors in the melanoma dataset. Top PCs accounting for 80% of the variance are highlighted in red.

(b) Same as in (a) but for the HNSCC dataset.

### Supplementary Figure 6

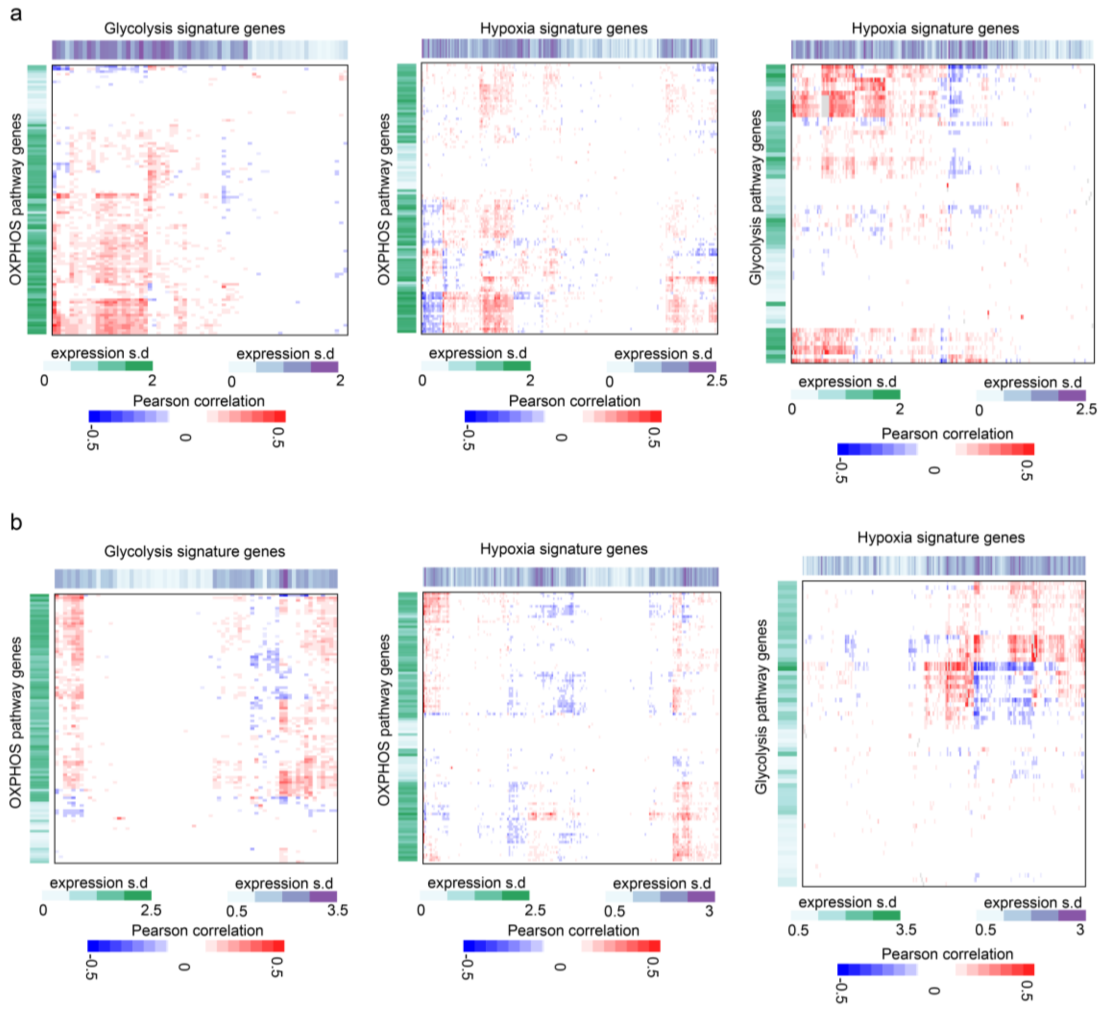


**Supplementary Figure 6. (Related to Figure 3)**

(a) Pairwise Pearson's correlation coefficients of gene expression levels between oxidative phosphorylation (OXPHOS), glycolysis, and response to hypoxia in malignant cells from the melanoma dataset. Color bars on the left and top show the standard deviations of expression levels of the genes.

(b) Same as in (a) but for HNSCC dataset.

**Supplementary Figure 7**

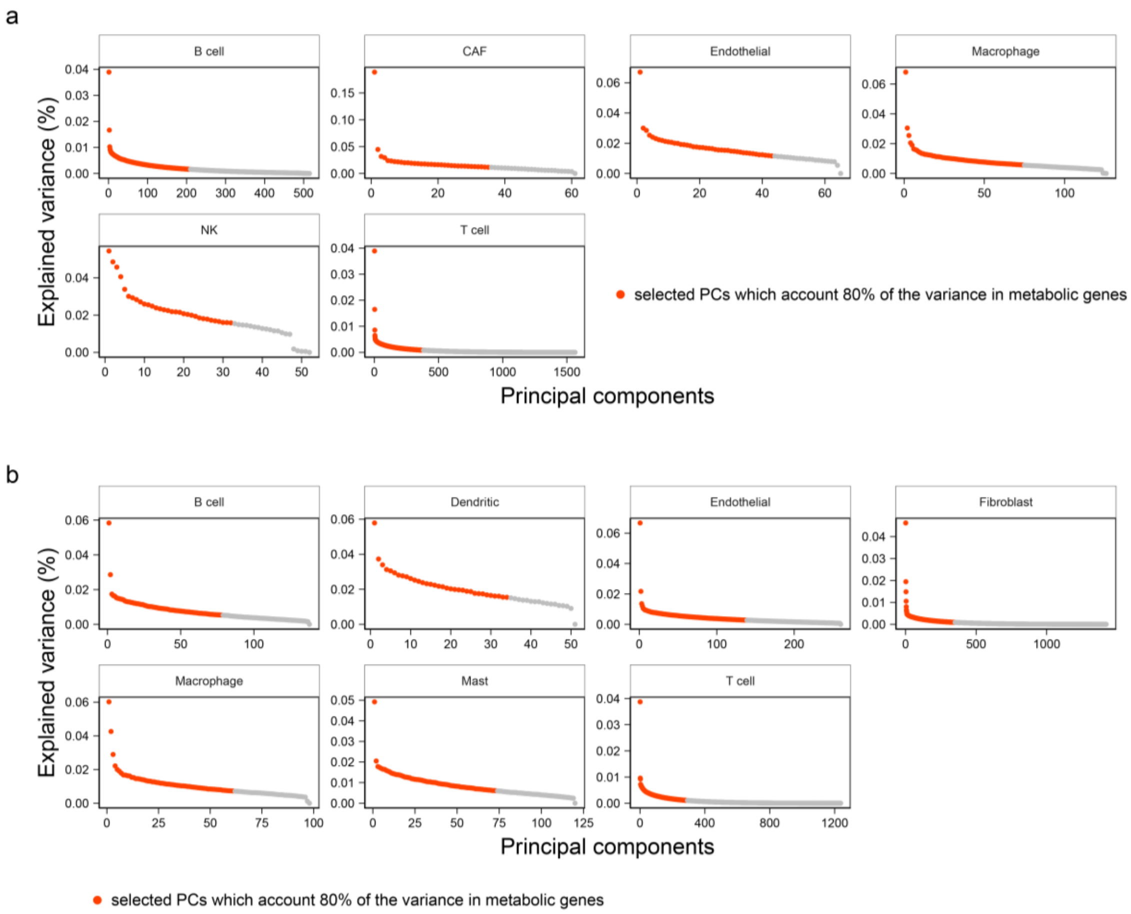


**Supplementary Figure 7. (Related to Figure 4)**

(a) Explained variance of principal components (PCs) from principal component analysis (PCA) of metabolic gene expression levels in different types of non-malignant cells from the melanoma dataset. Top PCs accounting for 80% of the variance are highlighted in red.

(b) Same as (a) but for HNSCC dataset.

**Table S1. Significantly enriched pathways in cancer associated fibroblasts (CAFs) compared to myofibroblasts in the HNSCC dataset**

| Pathway | NES[#] | p-value |
|---|---|---|
| Retinol metabolism | 1.600 | 0.000 |
| Arachidonic acid metabolism | 1.567 | 0.000 |
| Linoleic acid metabolism | 1.554 | 0.002 |
| Nitrogen metabolism | 1.538 | 0.001 |
| Galactose metabolism | 1.521 | 0.001 |
| Taurine and hypotaurine metabolism | 1.497 | 0.009 |
| N-Glycan biosynthesis | 1.477 | 0.005 |
| Other glycan degradation | 1.470 | 0.008 |
| Pentose and glucuronate interconversions | 1.469 | 0.006 |
| Amino sugar and nucleotide sugar metabolism | 1.413 | 0.008 |
| Glycosphingolipid biosynthesis - globo and isoglobo series | 1.390 | 0.040 |
| Glycosphingolipid biosynthesis - ganglio series | 1.378 | 0.038 |
| Glycosaminoglycan degradation | 1.377 | 0.030 |
| Sulfur metabolism | 1.366 | 0.053 |
| Metabolism of xenobiotics by cytochrome P450 | 1.362 | 0.019 |
| Arginine biosynthesis | 1.359 | 0.054 |
| Ascorbate and aldarate metabolism | 1.331 | 0.062 |
| Glycolysis / Gluconeogenesis | 1.318 | 0.049 |
| Ether lipid metabolism | 1.300 | 0.074 |
| Alanine, aspartate and glutamate metabolism | 1.294 | 0.081 |

[#] NES: Normalized enrichment scores
